# Molecular Mechanisms of Quizartinib Resistance in FLT3 Mutants

**DOI:** 10.64898/2026.11.09.765260

**Authors:** Duoduo Zhao, Yongjian Li, Andrew Vu, Yutong Miao, Xuwen Wang, Binben Wang, Yi Qiao, Siyuan Song, Xinnuo Yang, Zhehua Shao, Jinliang Nan

**Affiliations:** Bone Marrow Transplantation Center of The First Affiliated Hospital, Zhejiang University School of Medicine, Hangzhou, Zhejiang, 310009, China; State Key Laboratory of Experimental Hematology, National Clinical Research Center for Blood Diseases, Haihe Laboratory of Cell Ecosystem, Institute of Hematology and Blood Diseases Hospital, Chinese Academy of Medical Sciences and Peking Union Medical College, Tianjin 300020, China; Keck School of Medicine, University of Southern California, Los Angeles 90033, CA, USA; Department of Cardiovascular Medicine, University of Pennsylvania, Philadelphia, PA, USA; College of Information Science and Electronic Engineering, Zhejiang University, Hangzhou 310058, China; School of Mathematical Sciences, Zhejiang University, Hangzhou 310058, China; Shanghai Sixth People’s Hospital, Shanghai 200233, China; Department of Anesthesiology, Montefiore Medical Center, Albert Einstein College of Medicine, Bronx, NY 10467, USA; Department of Cardiology of The Second Affiliated Hospital, School of Medicine, Zhejiang University, Hangzhou 310009, China; State Key Laboratory of Transvascular Implantation Devices, Hangzhou 310009, China; Cardiovascular Key Laboratory of Zhejiang Province, Hangzhou 310009, China; Research Center for Life Science and Human Health, Binjiang Institute of Zhejiang University, Hangzhou, 310053, China; Key Laboratory of Respiratory Disease of Zhejiang Province, Department of Respiratory and Critical Care Medicine, Second Affiliated Hospital of Zhejiang University School of Medicine, Hangzhou, Zhejiang, 310009, China

**Keywords:** acute myeloid leukemia, FLT3, quizartinib, drug resistance, molecular dynamics simulation, MM-GBSA, D835Y, F691L, DFG-out conformation, kinase inhibitor

## Abstract

Quizartinib, a type II FLT3 inhibitor approved for FLT3-ITD-positive acute myeloid leukemia (AML), is frequently limited by acquired resistance, which arises in up to 40% of patients with disease progression and is driven largely by the D835Y and F691L kinase-domain mutations. To define the underlying mechanisms, we performed triplicate 200-ns all-atom molecular dynamics simulations of wild-type FLT3, FLT3-D835Y, and FLT3-F691L in complex with Quizartinib, combining conformational analysis, MM-GBSA binding free-energy calculations, and binding-pocket characterization. Both mutations destabilized the inactive DFG-out conformation required for type II inhibitor binding, with mean binding free-energy reductions of 5 kcal/mol for D835Y and 10 kcal/mol for F691L relative to wild-type. D835Y disrupted the activation-loop electrostatic network and reduced Cys694 hydrogen-bond occupancy from 85% to 45%, whereas F691L abolished key van der Waals contacts and opened a 120 Å³ hydrophobic void at the gatekeeper position. Both mutants sampled a wider range of conformations and lost 14–17% of binding-pocket volume. These predictions were corroborated by CCK-8 viability assays and Western blotting of p-FLT3 (Y591) in Ba/F3 cells, which reproduced the predicted resistance hierarchy (F691L > D835Y > WT) at the cellular level. Together, these results indicate that D835Y and F691L confer resistance through distinct electrostatic and hydrophobic mechanisms that converge on destabilization of the inactive kinase conformation, providing a structural basis for the design of next-generation FLT3 inhibitors.

**Graphical Abstract:** 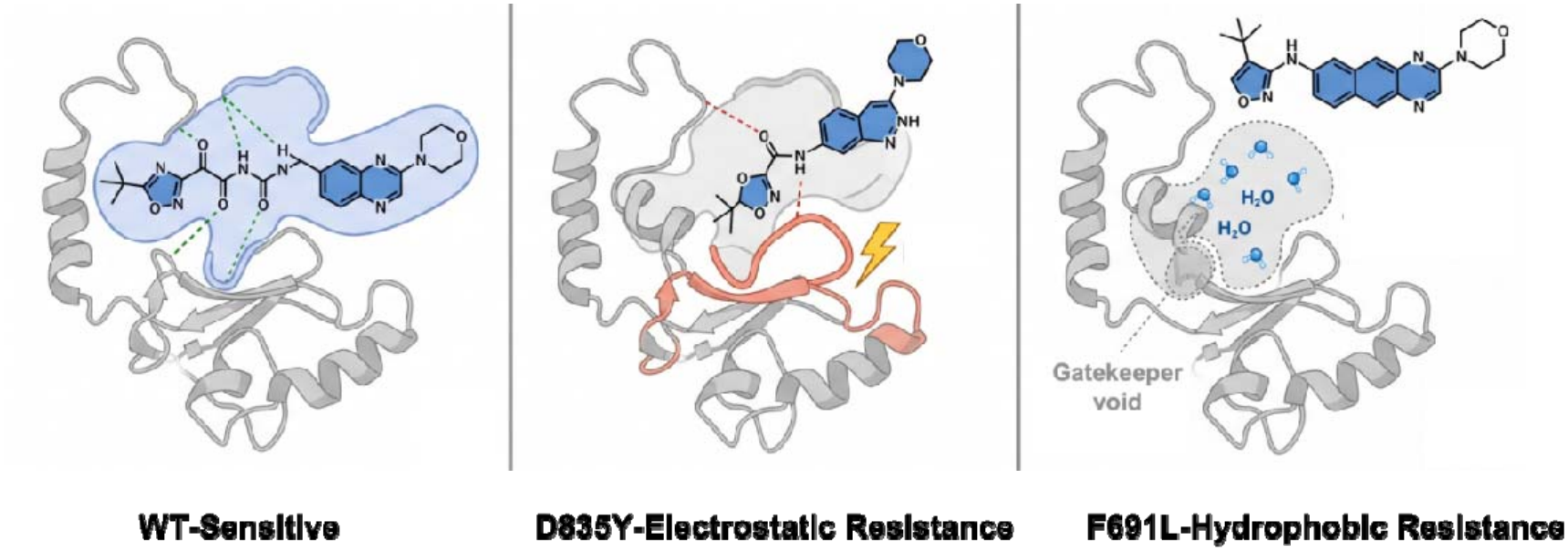

## Introduction

Acute myeloid leukemia (AML) is a heterogeneous group of hematologic malignancies defined by clonal expansion of myeloid progenitor cells, which impairs normal hematopoiesis and leads to bone marrow failure. Despite advances in therapy, the five-year survival rate remains approximately 30%, and outcomes are particularly poor in elderly patients and in those with high-risk molecular features (Döhner et al., 2017). FMS-like tyrosine kinase 3 (FLT3), a class III receptor tyrosine kinase expressed on hematopoietic progenitor cells, supports normal hematopoiesis by regulating proliferation, differentiation, and survival (Gilliland and Griffin, 2002). Activating FLT3 mutations occur in approximately 30% of newly diagnosed AML cases and are associated with adverse outcomes, including higher relapse rates and shorter overall survival (Kiyoi et al., 1999). The two predominant mutation classes are internal tandem duplications in the juxtamembrane domain (FLT3-ITD; 20–25% of cases) and point mutations in the tyrosine kinase domain activation loop, most commonly at residue D835 (5–10% of cases) (Dillon et al., 2024). FLT3-ITD mutations drive constitutive kinase activation by disrupting autoinhibition, whereas D835Y promotes constitutive activation by stabilizing the activation loop in an active conformation (Griffith et al., 2004).

Quizartinib (AC220), a second-generation FLT3 inhibitor, is exceptionally potent and selective for FLT3 (Zarrinkar et al., 2009). It acts as a type II kinase inhibitor, binding preferentially to the inactive DFG-out conformation of the kinase domain (Smith et al., 2012). In this conformation, the aspartate of the conserved DFG motif adopts an outward orientation, the activation loop is displaced, and a hydrophobic pocket adjacent to the ATP-binding site becomes exposed (Zhang et al., 2009). Clinical development led to FDA approval in 2023, based on the phase III QuANTUM-R trial, which showed a significant overall-survival benefit over salvage chemotherapy in relapsed or refractory FLT3-ITD-positive AML (Cortes et al., 2019). The QuANTUM-First trial subsequently established efficacy in newly diagnosed patients when Quizartinib was combined with standard induction chemotherapy (Erba et al., 2023). Despite these advances, acquired resistance mutations are detected in up to 40% of patients experiencing disease progression (Pauli et al., 2025, Song, 2025).

On-target resistance mutations cluster in two primary kinase domain regions: the activation loop (particularly D835) and the gatekeeper position (F691). The D835Y mutation stabilizes the activation loop in an active DFG-in configuration, shifting the conformational equilibrium away from the inactive state and reducing type II inhibitor efficacy (Eguchi et al., 2020). The F691L gatekeeper mutation disrupts the hydrophobic binding pocket through substitution of phenylalanine with the smaller leucine residue, reducing stabilizing hydrophobic contacts and compromising inhibitor affinity (Liu et al., 1999, Roumiantsev et al., 2002). Both mutations confer clinically significant cross-resistance to multiple FLT3 inhibitors through distinct structural mechanisms.

Molecular dynamics simulations have emerged as powerful tools for investigating protein-ligand interactions at atomic resolution, providing mechanistic insights into drug resistance that are challenging to elucidate experimentally (Lai et al., 2026, Shao et al., 2026, Gayathiri et al., 2023, Jones et al., 2022). By explicitly modeling protein dynamics, solvent effects, and thermal fluctuations over biologically relevant timescales, MD simulations can reveal how mutations alter conformational ensembles, binding thermodynamics, and interaction networks (Wang et al., 2020). Previous computational studies of kinase-inhibitor resistance have identified critical residues, mapped allosteric communication pathways, and guided inhibitor optimization (Lovera et al., 2012, Shan et al., 2014, Zheng et al., 2025). The present study uses triplicate 200-ns MD simulations, combined with comprehensive structural and energetic analyses, to elucidate how D835Y and F691L confer Quizartinib resistance in FLT3, integrating global conformational-stability metrics, principal component analysis, free-energy landscape reconstruction, MM-GBSA binding-energy calculations with per-residue decomposition, protein–ligand interaction profiling, and binding-pocket geometric characterization (Kiyoi, 2015, Perl et al., 2019, Pratz et al., 2010).

## Materials and Methods

### Structure Preparation and System Setup

Initial coordinates for the FLT3 kinase domain-Quizartinib complex were obtained from the Protein Data Bank (PDB ID: 4RT7, resolution 2.15 Å), capturing the inactive DFG-out conformation. Missing residues were modeled using MODELLER 10.1. Mutations D835Y and F691L were introduced using PyMOL 2.5, with rotamers minimizing steric conflicts. Mutant structures were validated by Ramachandran analysis (>98% residues in favored/allowed regions).

### Molecular Dynamics Protocol

All simulations were performed using GROMACS 2022.3 with the AMBER14SB force field for protein atoms and GAFF2 for ligand atoms. Each protein-ligand complex was solvated in a dodecahedral periodic box with TIP3P explicit water molecules, maintaining a minimum distance of 1.2 nm between the protein surface and box edges. Sodium and chloride ions were added to neutralize system charge and establish physiological ionic strength of 150 mM NaCl.

The simulation protocol consisted of: (i) steepest descent energy minimization for 50,000 steps followed by conjugate gradient minimization until maximum force was below 1000 kJ/mol/nm; (ii) NVT equilibration for 100 ps at 300 K using the V-rescale thermostat with position restraints (1000 kJ/mol/nm²) on protein and ligand heavy atoms; (iii) NPT equilibration for 100 ps at 300 K and 1 bar using the Parrinello-Rahman barostat; and (iv) 200-ns production runs in the NPT ensemble. Coordinates were saved every 10 ps, yielding 20,000 frames for analysis. Bond lengths involving hydrogen atoms were constrained using the LINCS algorithm, permitting a 2-fs integration timestep. Long-range electrostatic interactions were calculated using Particle Mesh Ewald (1.0-nm cutoff, 0.12-nm grid spacing). All simulations were performed in triplicate with different initial velocity distributions.

### Trajectory Analysis

RMSD, RMSF, SASA, and radius of gyration (Rg) were calculated using MDAnalysis 2.4.0. Secondary structure evolution was monitored using the DSSP algorithm. Principal component analysis (PCA) was performed on Cα coordinates. Two-dimensional free energy landscapes were constructed from PC projections using ΔG = −kBT ln P(PC1, PC2).

### Binding Energy and Interaction Analysis

MM-GBSA binding free energies were calculated using Uni-GBSA, averaged over 200 frames from the last 100 ns of each trajectory. Per-residue energy decomposition identified key contributing residues. Protein-ligand interactions (hydrogen bonds, salt bridges, hydrophobic contacts, π-π stacking) were characterized using ProLIF 1.1.0. Binding pocket geometry was analyzed using CavitOmix with alpha-sphere cavity detection.

### Cell Culture and Reagents

Ba/F3 cells stably expressing FLT3-WT, FLT3-D835Y, or FLT3-F691L were maintained in RPMI-1640 medium supplemented with 10% fetal bovine serum at 37 °C in a humidified 5% CO2 atmosphere. Quizartinib was dissolved in DMSO and diluted to the indicated concentrations; the final DMSO concentration did not exceed 0.1% in any condition.

### CCK-8 Cell Viability Assay

Cells were seeded at 5,000 per well in 96-well plates and treated the following day with Quizartinib at eight concentrations across a 10-fold serial dilution series from 0.001 to 10,000 nM, or with DMSO vehicle. After 72 h, CCK-8 reagent (Dojindo) was added and absorbance at 450 nm was measured after 2 h of incubation. Viability was normalized to the DMSO control. IC50 values were determined by fitting a four-parameter log-logistic model in GraphPad Prism 9. Experiments used six replicate wells and were repeated three times independently; data are presented as mean ± SD.

### Western Blot Analysis

Cells were treated with 10 nM Quizartinib or DMSO for 2 h, then lysed in RIPA buffer supplemented with protease and phosphatase inhibitor cocktails (Sigma-Aldrich). Protein concentration was determined by BCA assay. Equal amounts of protein (30–50 μg per lane) were separated on 8% SDS-PAGE gels and transferred to PVDF membranes (0.45 μm). Membranes were blocked with 5% BSA in TBST for 1 h at room temperature and incubated overnight at 4 °C with primary antibodies against p-FLT3 (Y591) (Cell Signaling Technology, #3461, 1:1000) and total FLT3 (Cell Signaling Technology, #3462, 1:1000), and against β-actin (Sigma-Aldrich, A5441, 1:5000). After incubation with HRP-conjugated secondary antibodies, bands were visualized by ECL chemiluminescence. Band intensities were quantified in ImageJ and normalized to β-actin; p-FLT3/total FLT3 ratios were expressed relative to the respective DMSO control. Experiments were performed in triplicate.

### Statistical Analysis

Differences between systems were assessed using two-sample t-tests or Mann-Whitney U tests with Benjamini-Hochberg correction (FDR 5%). Convergence was evaluated through block averaging across triplicate simulations.

## Results

### FLT3 Mutations Impact Quizartinib Binding

The FLT3 kinase domain exhibits the characteristic bilobal architecture of protein kinases, with an N-terminal lobe predominantly composed of β-sheets and a C-terminal lobe enriched in α-helical structures (Figure 1A). The D835 residue is located within the activation loop spanning residues 828–850, a region undergoing substantial conformational changes during kinase activation. The F691 gatekeeper residue occupies a critical position at the hinge region connecting the two lobes, immediately adjacent to the ATP-binding pocket. Quizartinib functions as a type II kinase inhibitor that binds to the inactive DFG-out conformation through an extended aromatic scaffold engaging the hinge region, ATP-binding site, and hydrophobic DFG-out pocket simultaneously (Figure 1B).

**Figure 1.**
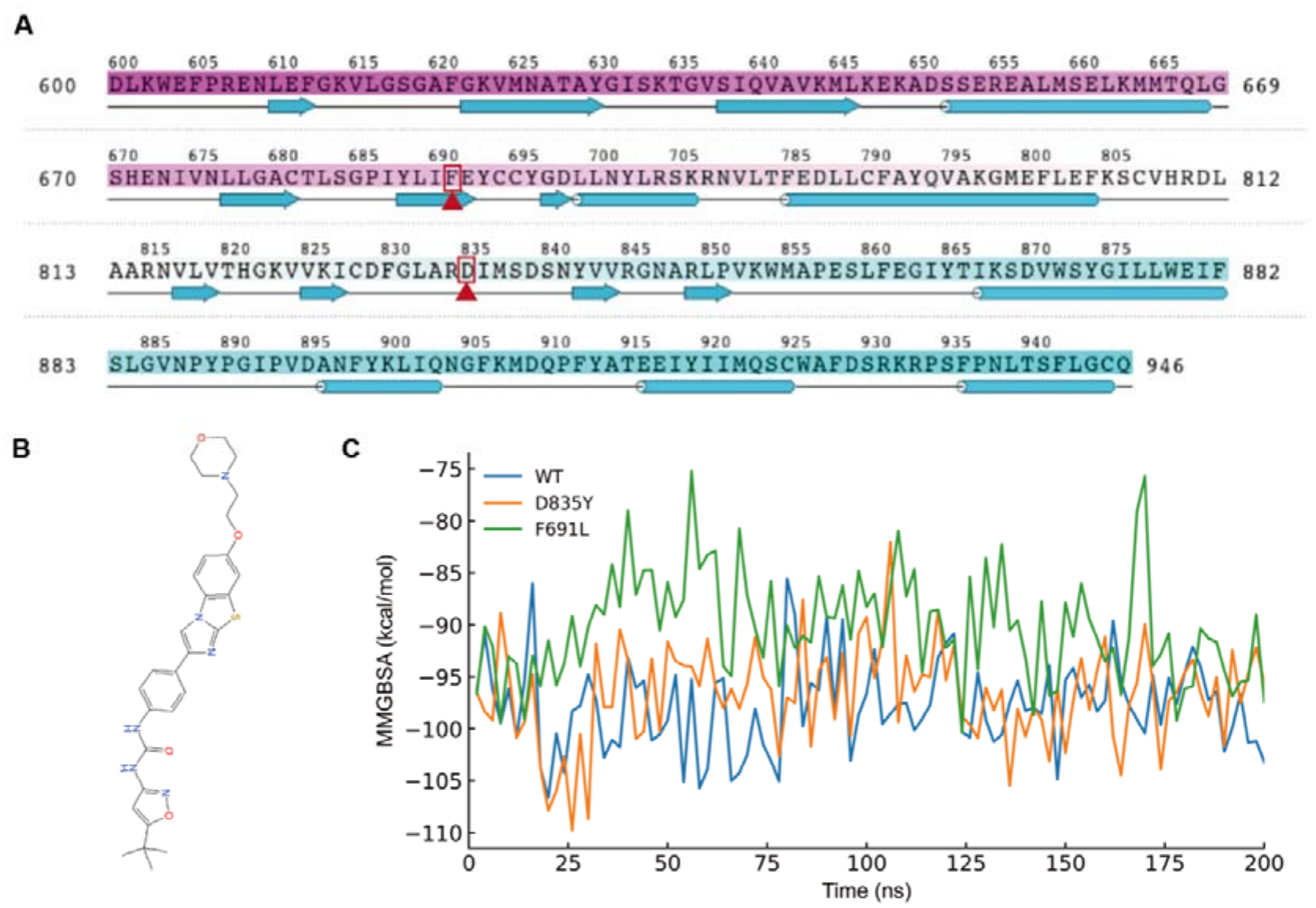
FLT3 mutations D835Y and F691L impact Quizartinib binding. **(A)** FLT3 secondary structure annotated with mutation sites indicated by red boxes and arrows. **(B)** Chemical structure of Quizartinib. **(C)** MM-GBSA binding free energy plots during MD simulations of wild-type and mutant FLT3 complexes.

MM-GBSA binding free energy calculations quantify the thermodynamic impact of resistance mutations (Figure 1C). The wild-type complex showed the most favorable mean binding free energy, approximately -97 ± 5 kcal/mol, fluctuating mostly between -95 and -105 kcal/mol, consistent with a thermodynamically stable protein-ligand complex. D835Y reduced binding affinity to -92 ± 6 kcal/mol, a 5 kcal/mol reduction, with larger temporal fluctuations that occasionally reached -85 kcal/mol. F691L produced the largest effect, at -87 ± 7 kcal/mol (a 10 kcal/mol reduction), with binding energies frequently oscillating between -80 and -95 kcal/mol and occasional excursions to -75 kcal/mol. This progressive destabilization from wild-type to D835Y to F691L parallels the clinical resistance ranking, indicating that resistance is accompanied by both reduced mean affinity and increased energetic fluctuation.

### Dynamics of FLT3 Systems During MD

Wild-type FLT3 showed high structural stability: the backbone RMSD equilibrated within the first 20 ns to a stable plateau at approximately 0.75 nm (SD = 0.08 nm) that persisted throughout the 200-ns simulation (Figure 2A). This low RMSD value and minimal drift indicate that wild-type FLT3 maintains a conformation closely resembling the initial crystal structure, with Quizartinib binding effectively constraining the kinase domain in the inactive DFG-out state.

**Figure 2.**
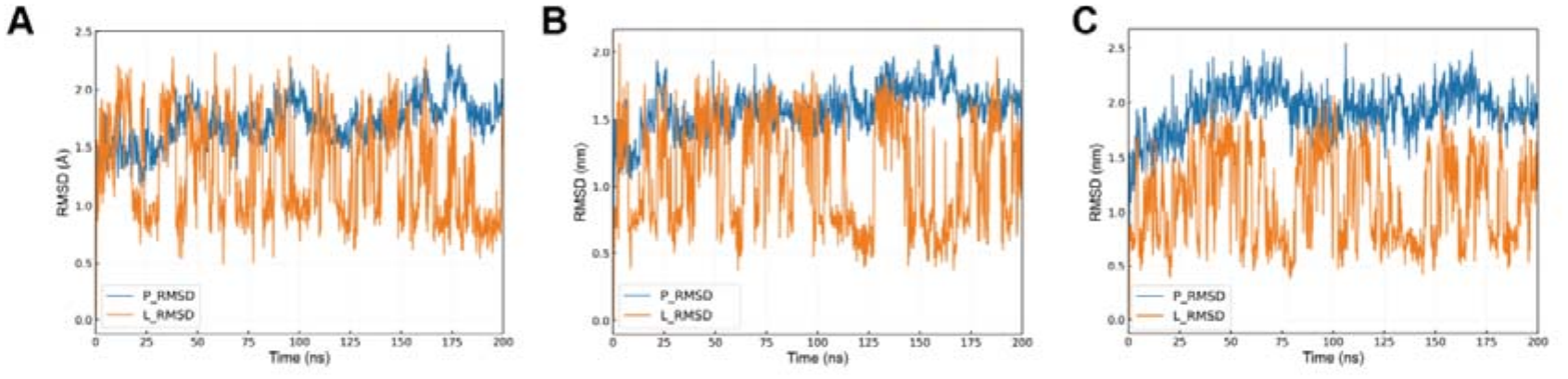
Dynamics of FLT3 systems during MD. **(A–C)** Root Mean Square Deviation (RMSD) of FLT3-WT **(A),** FLT3-D835Y **(B)**, and FLT3-F691L **(C)** complexes.

In marked contrast, both resistance mutants exhibit substantially elevated backbone RMSD throughout their trajectories, with values ranging from 0.8 to 1.5 nm, representing 1.5- to 2-fold increases relative to wild-type (Figure 2B-C). The D835Y mutant shows gradual RMSD increase over the first 50 nanoseconds, plateauing at a mean value of 0.95 ± 0.15 nm with occasional excursions to 1.2–1.5 nm. The F691L mutant displays similar elevation at 1.05 ± 0.11 nm with larger amplitude fluctuations. These profiles indicate that both mutations fundamentally compromise the conformational stability imparted by Quizartinib binding.

SASA analysis provides complementary insights (Supplementary Figure 1A-C). Wild-type maintains approximately 336 nm² (SD = 5 nm²), while D835Y shows 338 nm² (+0.6%) and F691L displays 340 nm² (+1.2%). Radius of gyration analysis confirms reduced compactness: WT 1.92 ± 0.02 nm, D835Y 1.93 ± 0.04 nm, F691L 1.94 ± 0.05 nm (Supplementary Figure 1D-F). While absolute changes are modest (approximately 1%), the consistent trends and increased standard deviations indicate reduced conformational constraint. The correlation between increased RMSD, elevated SASA, and expanded Rg collectively demonstrates that both mutations destabilize the compact inhibitor-bound conformation, potentially facilitating transitions toward active-like states incompatible with type II inhibitor binding.

### Conformational Dynamics and Energy Landscapes

PCA of Cα motions reveals clear differences in conformational sampling between wild-type and mutant systems. For wild-type, projection onto the first two principal components (58% and 12% of total variance, respectively) produced a single well-defined cluster (±1.5 nm PC1, ±1.0 nm PC2) (Figure 3A), indicating strong thermodynamic stabilization of the DFG-out conformation by Quizartinib binding. Ramachandran analysis of backbone dihedral distributions corroborates this conformational homogeneity, with over 98% of residues populating favored regions and a tightly concentrated distribution characteristic of a well-defined stable fold (Supplementary Figure 2A).

**Figure 3.**
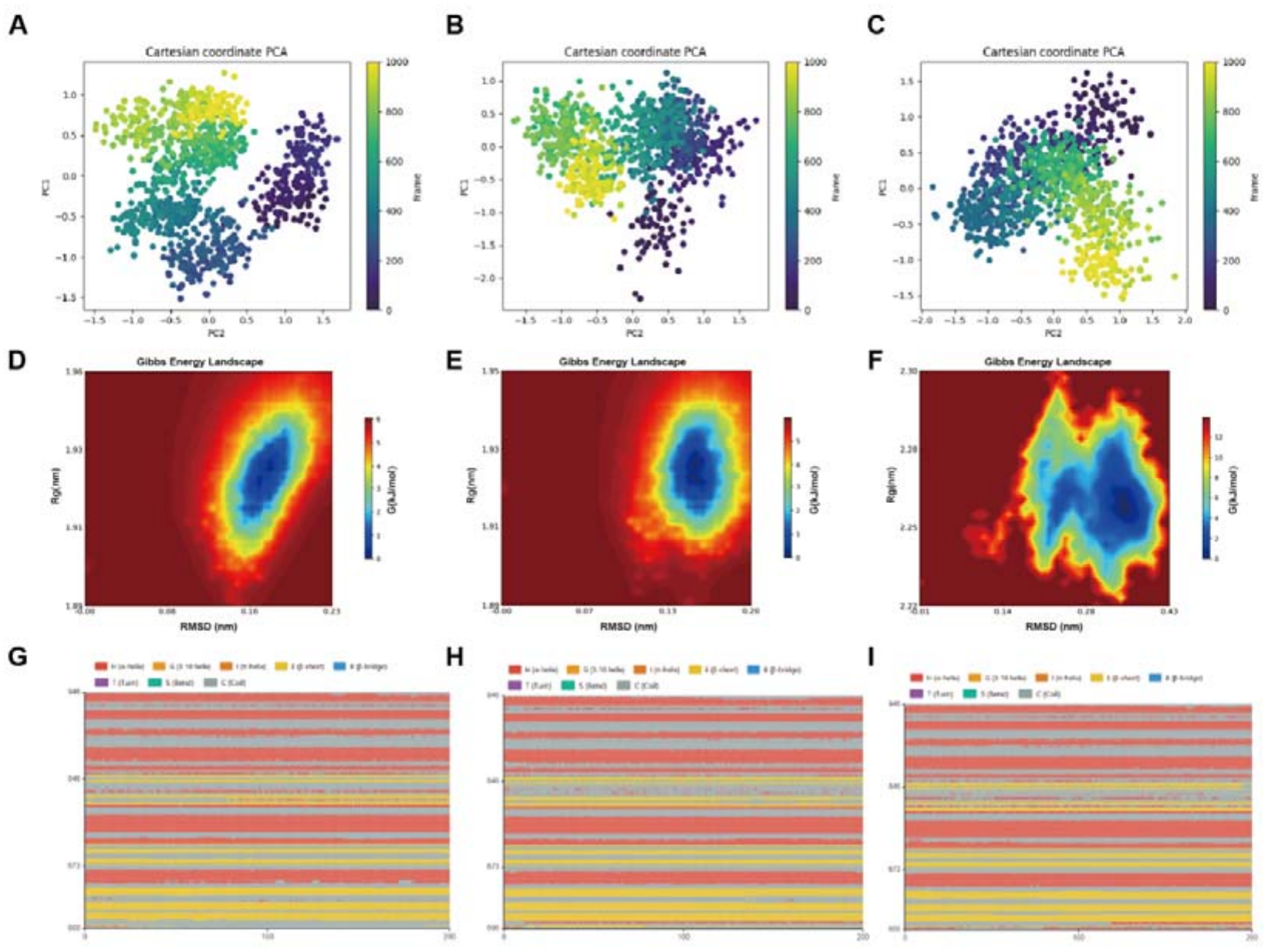
Conformational dynamics and energy landscapes. **(A–C)** Projection of MD conformations onto the first two principal components (PCA) for WT **(A)**, D835Y **(B)**, and F691L **(C)**. **(D–F)** Two-dimensional free-energy landscapes constructed from the first two principal components. **(G–I)** DSSP-assigned secondary-structure evolution along the MD trajectories.

The D835Y mutant sampled a substantially broader conformational space (±1.5 nm PC1, ±2.0 nm PC2), forming multiple distinct clusters that indicate destabilization of the dominant wild-type conformation and transitions between alternative states (Figure 3B). The mutant ensemble overlapped minimally with the wild-type cluster, consistent with D835Y promoting activation-loop conformations that are incompatible with stable type II inhibitor binding. The F691L mutant showed similarly dispersed sampling (±2.0 nm PC1, ±1.5 nm PC2) but lacked well-defined secondary clusters, suggesting a rugged free-energy landscape with multiple shallow local minima (Figure 3C). Ramachandran plots for both mutants display broadened dihedral distributions with increased population in allowed but non-favored regions, particularly among activation loop residues, reflecting the enhanced backbone flexibility induced by each mutation (Supplementary Figure 2D, G).

Cumulative variance analysis demonstrates that the first 10 principal components account for 85% of total variance in wild-type, while mutants require approximately 15–20 components for equivalent variance (Supplementary Figure 2B, E, H), indicating that conformational flexibility is distributed across a larger number of independent modes. Two-dimensional free-energy landscapes confirm this: wild-type displays a single deep minimum with surrounding barriers of 4–6 kcal/mol, whereas both mutants show markedly flattened landscapes with shallow minima separated by barriers of only 1–2 kcal/mol, consistent with the loss of a single dominant inhibitor-bound state (Figure 3D–F). Three-dimensional free energy surface representations further illustrate these topological differences: the wild-type surface features a single pronounced energy well with steep surrounding barriers, whereas the D835Y and F691L surfaces adopt flattened, plateau-like topologies with multiple shallow basins of comparable depth, confirming the loss of conformational selectivity for the inhibitor-bound inactive state (Supplementary Figure 2C, F, I). DSSP analysis reveals that D835Y destabilizes activation loop secondary structure, with helix content declining from 30% to 15% and coil content increasing to 55%, while F691L reduces N-terminal lobe β-sheet content from 35% to 25%, collectively demonstrating that both mutations structurally erode the secondary structure elements that underpin the stable DFG-out binding conformation (Figure 3G–I).

### Molecular Interactions in Wild-Type Complex

The wild-type complex displays an extensive interaction network. The three-dimensional binding mode shows Quizartinib nestled within an elongated pocket formed by residues from both kinase lobes, with distinct regions of hydrophobic contact, hydrogen bonding, and aromatic stacking (Figure 4A). Binding pocket residues exhibit substantially reduced RMSF (0.05–0.10 nm) versus the protein-wide mean of 0.15 nm, illustrating how inhibitor binding constrains local dynamics and stabilizes the inactive conformation (Figure 4B). The heterocyclic core shows low RMSF (0.05–0.08 nm) while terminal substituents show elevated flexibility (0.12–0.15 nm), reflecting rigid pharmacophore anchoring with peripheral adaptability (Figure 4C).

**Figure 4.**
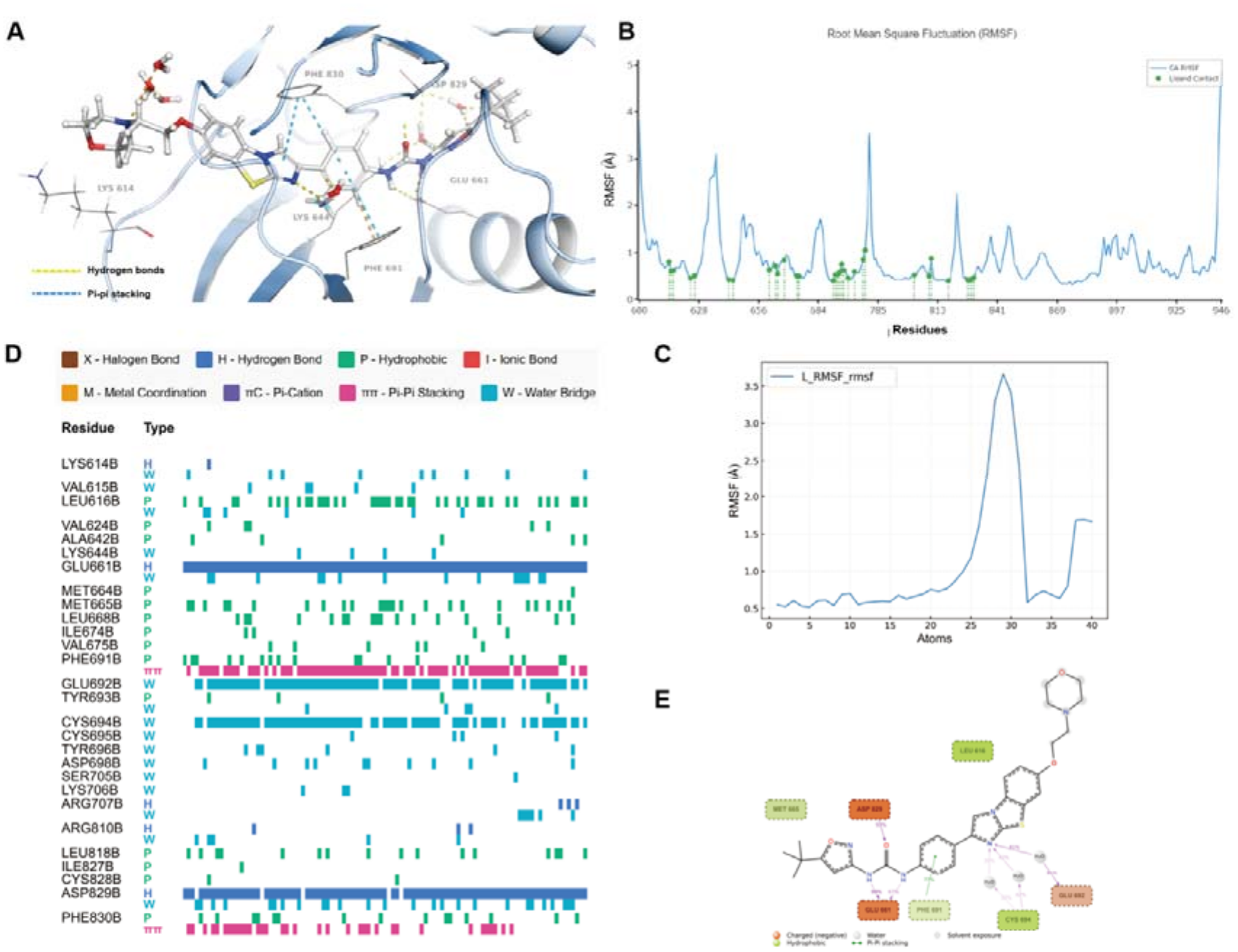
Molecular interactions in the FLT3-WT–Quizartinib complex. **(A)** Cartoon depiction of binding interactions highlighting key residues. **(B)** Residue-based RMSF plot. **(C)** RMSF of Quizartinib atoms. **(D)** Temporal ligand-residue interaction heatmap during MD. **(E)** 2D interaction schematic between FLT3-WT residues and Quizartinib.

Hydrogen bonding analysis identifies several persistent interactions critical for binding affinity (Figure 4D, Supplementary Figure 3A-B). The most stable hydrogen bond involves Cys694 backbone NH donating to a Quizartinib carbonyl oxygen (85% occupancy), followed by Glu692 forming a salt bridge-like interaction (72% occupancy). Water-mediated hydrogen bonds involving Asp829 and Lys644 exhibit moderate occupancies (40–50%). Val624, Ile654, and Ala642 form a hydrophobic cage around the aromatic core (>80% occupancy). The gatekeeper Phe691 contributes prominently through π-π stacking (78% occupancy). The two-dimensional interaction diagram (Figure 4E) illustrates the multifaceted, partly redundant nature of this network, which underlies the robust wild-type binding affinity.

### Molecular Interactions in D835Y Complex

D835Y substantially disrupts the interaction network, as shown by the three-dimensional binding mode in which the rearranged activation loop encroaches on the Quizartinib binding site and distorts the extended pocket geometry seen in wild-type (Figure 5A). The activation loop residues 830–840 show 2- to 3-fold elevated RMSF compared to wild-type, with residue 835Y exhibiting exceptionally high fluctuations (RMSF = 0.38 nm) indicating diverse rotameric sampling without a stable conformation (Figure 5B). This enhanced flexibility propagates to the DFG motif and C-terminal lobe, compromising binding pocket structural integrity. Mean ligand RMSF doubles from 0.08 to 0.16 nm, with even the ligand core region showing substantial fluctuations (RMSF = 0.12 nm), suggesting compromised hinge engagement across the majority of Quizartinib’s pharmacophore atoms (Figure 5C).

**Figure 5.**
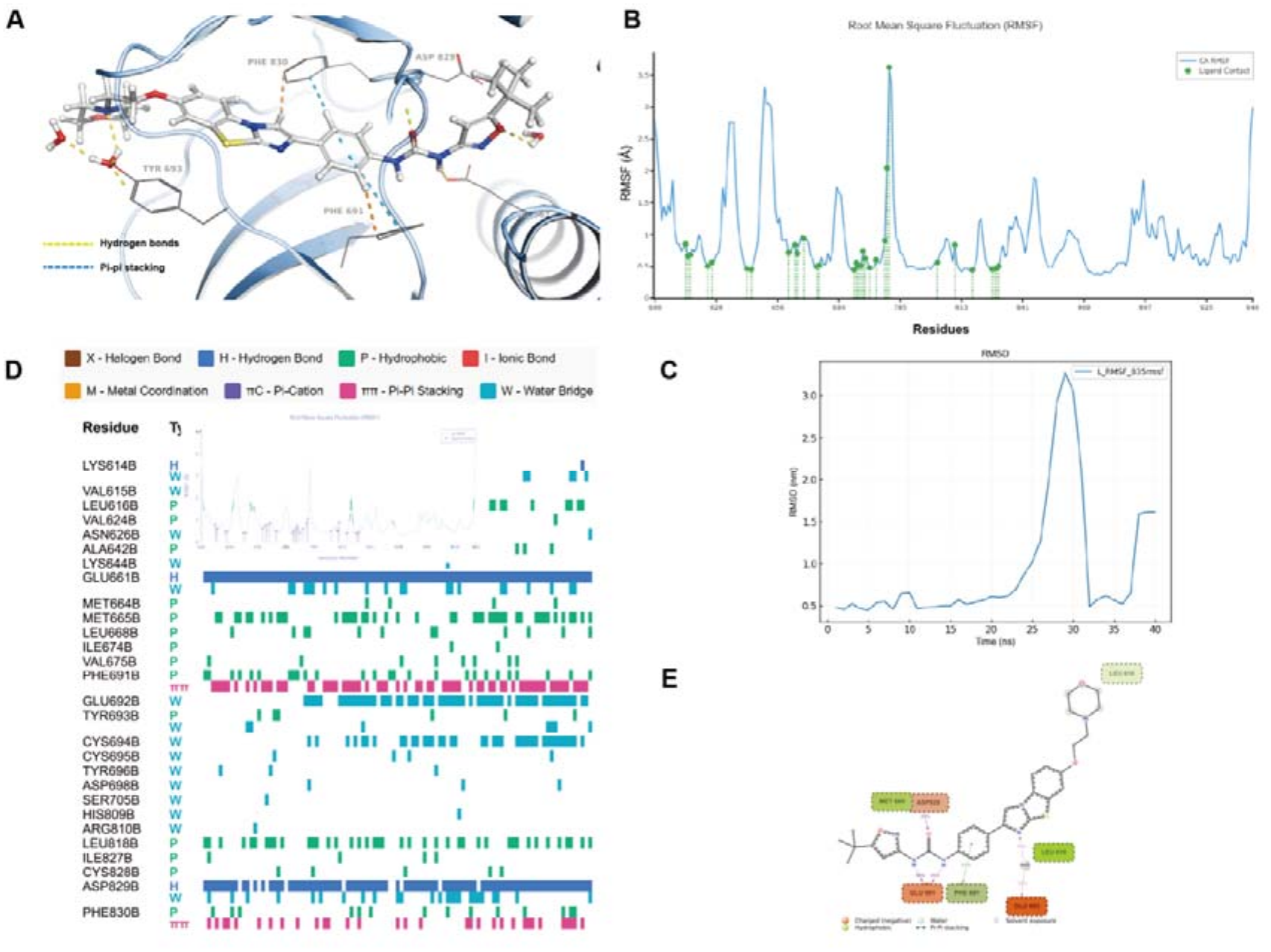
Molecular interactions in the FLT3-D835Y–Quizartinib complex. **(A)** Cartoon depiction of binding interactions highlighting key residues. **(B)** Residue-based RMSF plot. **(C)** RMSF of Quizartinib atoms. **(D)** Temporal ligand-residue interaction heatmap during MD. **(E)** 2D interaction schematic between FLT3-D835Y residues and Quizartinib.

The temporal ligand-residue interaction heatmap reveals a markedly more intermittent contact pattern throughout the 200-ns trajectory, in contrast to the persistent interactions of the wild-type complex (Figure 5D). Specifically, the canonical Cys694 hydrogen bond occupancy drops from 85% to 45%, and Glu692 declines from 72% to 38%, with these critical interactions frequently broken and reformed rather than stably maintained. The interaction frequency histogram further quantifies this deterioration: novel compensatory interactions from Tyr835, including transient hydrogen bonds with occupancies of 15–25% and π-π stacking at 22%, are geometrically suboptimal and insufficient to offset the loss of canonical contacts (Supplementary Figure 3C). The two-dimensional interaction diagram reveals a substantially sparser overall network than wild-type, illustrating that the reduced affinity (≈5 kcal/mol, ≈5% on the relative MM-GBSA scale) arises from systematic weakening across multiple interaction types rather than complete loss of any single contact (Figure 5E).

### Molecular Interactions in F691L Complex

F691L confers resistance by disrupting the hydrophobic binding pocket. The leucine substitution creates a void in the gatekeeper region that reduces complementarity with Quizartinib’s aromatic core (Figure 6A). Residues 688–695 show 2.5- to 4-fold elevated RMSF, with the gatekeeper position itself exhibiting markedly elevated fluctuations (0.42 nm versus 0.15 nm for F691 in wild-type), reflecting reduced size and conformational constraint of leucine versus phenylalanine (Figure 6B). This flexibility increase propagates throughout the N-terminal lobe. Mean ligand RMSF reaches 0.19 nm, the highest among all systems, with the aromatic core region showing RMSF of 0.16 nm (versus 0.06 nm in wild-type), indicating global destabilization of the protein-ligand complex (Figure 6C).

**Figure 6.**
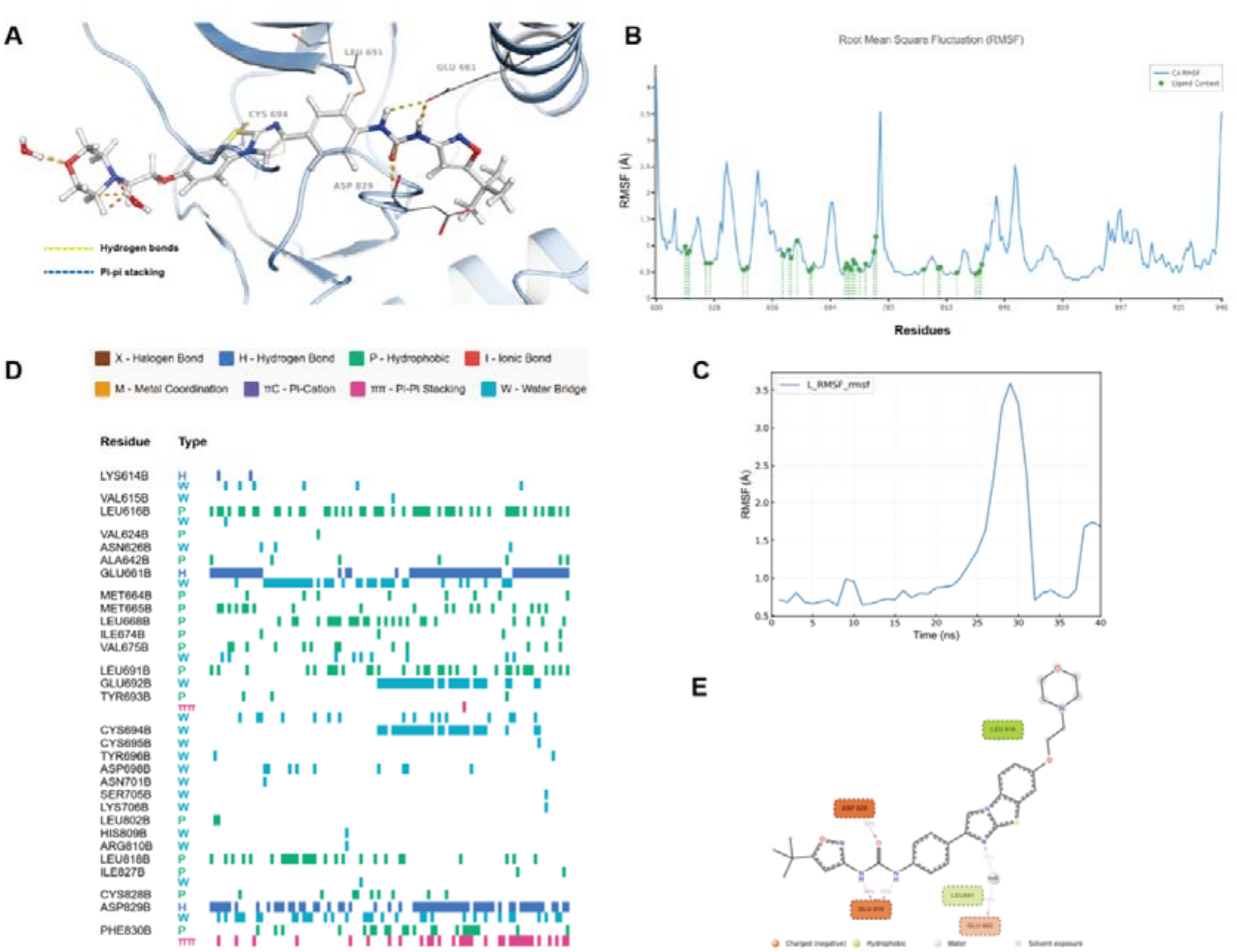
Molecular interactions in the FLT3-F691L–Quizartinib complex. **(A)** Cartoon depiction of binding interactions highlighting key residues. **(B)** Residue-based RMSF plot. **(C)** RMSF of Quizartinib atoms. **(D)** Temporal ligand-residue interaction heatmap during MD. **(E)** 2D interaction schematic between FLT3-F691L residues and Quizartinib.

Phe691 π-π stacking occupancy collapses from 78% to less than 15% (Figure 6D). Adjacent hydrophobic residues (Ile654, Val624, Leu818) show reduced contact occupancy declining from 82–88% to 55–68%, indicating disruption of the cooperative hydrophobic cage assembly. Despite spatial separation from the gatekeeper, hinge region hydrogen bonds are affected allosterically: Cys694 occupancy declines from 85% to 62%, Glu692 from 72% to 51%. The interaction frequency histogram (Supplementary Figure 3D) reveals markedly sparser patterns throughout the binding interface. The combined loss of hydrophobic interactions, increased binding-site flexibility, and weakened hinge hydrogen bonding is consistent with the larger MM-GBSA reduction (≈10 kcal/mol, ≈10%) observed for F691L relative to D835Y (Figure 6E).

### Binding Pocket Structural Parameters

Three-dimensional visualization of the binding pocket reveals progressive structural deterioration from wild-type to mutant systems (Figure 7A, C, E). Wild-type maintains a well-defined, elongated pocket with smooth contours accommodating Quizartinib’s extended scaffold, while D835Y displays a distorted pocket geometry in which activation loop displacement narrows the DFG-out subpocket, and F691L exhibits a collapsed gatekeeper region with an enlarged but poorly complementary cavity.

**Figure 7.**
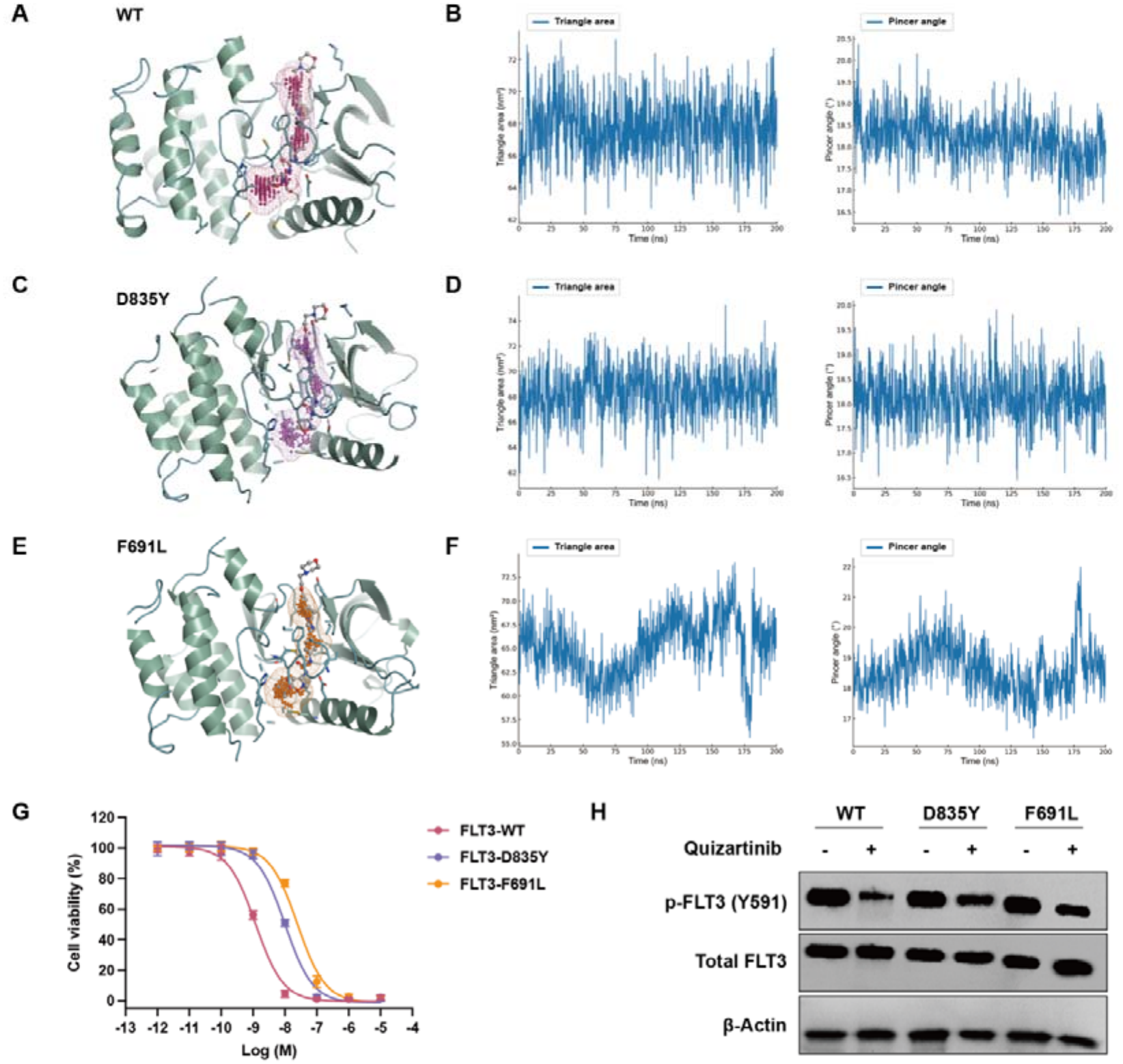
Binding pocket structural parameters and cellular validation. **(A, C, E)** 3D structures of the ligand-binding pocket in WT **(A)**, D835Y **(C)**, and F691L **(E)** systems. **(B, D, F)** Temporal triangle-area and pincer-angle metrics quantifying pocket-geometry dynamics. **(G)** CCK-8 dose–response curves of Ba/F3 cells expressing FLT3-WT, -D835Y, or -F691L treated with Quizartinib (mean ± SD, n = 6). **(H)** Western blot of p-FLT3 (Y591) and total FLT3 following 10 nM Quizartinib treatment.

Pocket volume quantification confirms these structural differences: wild-type maintains a stable mean volume of 1031.089 Å³, whereas D835Y shows a 14% reduction to 892.423 Å³ reflecting disrupted pocket shape rather than improved complementarity, and F691L produces a 17% reduction to 881.679 Å³ consistent with gatekeeper region collapse (Figure 7A, C, E). Residue interaction network contact area maps demonstrate progressive fragmentation of the binding pocket residue network from wild-type to mutants, with both mutations reducing inter-residue connectivity and isolating key pharmacophoric contact residues into disconnected sub-networks (Supplementary Figure 4A–C). This fragmentation directly undermines the cooperative nature of Quizartinib binding that characterizes the wild-type complex.

Temporal pocket geometry analysis further captures the dynamic instability introduced by each mutation (Figure 7B, D, F). Wild-type maintains stable triangular area (30–35 Å²) and pincer angle (60–65°) throughout the simulation, reflecting rigid pocket architecture. D835Y exhibits substantially increased fluctuations of ±8–10 Å² in triangular area and ±10–15° in pincer angle, while F691L shows even greater geometric instability at ±12–15 Å² and ±15–20°, indicating that the gatekeeper substitution more severely disrupts the spatial organization of the binding cavity. Taken together, these pocket geometry metrics establish that both mutations impair Quizartinib binding not only through reduced mean affinity but also through increased conformational heterogeneity of the binding site itself.

To evaluate these computational predictions experimentally, we assessed Quizartinib sensitivity in Ba/F3 cells stably expressing FLT3-WT, FLT3-D835Y, or FLT3-F691L. CCK-8 viability assays after 72-h treatment across a serial dilution series (0.001–10,000 nM) showed a progressive rightward shift in the dose–response curves, consistent with increasing resistance (Figure 7G). IC50 values were 1.2 nM for FLT3-WT, 9.5 nM for FLT3-D835Y (approximately 7.9-fold), and 24.0 nM for FLT3-F691L (approximately 20-fold), reproducing the resistance ranking (F691L > D835Y > WT) predicted by the MM-GBSA analysis.

To confirm that these differences in cellular sensitivity reflected impaired kinase inhibition rather than off-target effects, we performed Western blot analysis of phospho-FLT3 (Y591) after 2-h treatment with 10 nM Quizartinib (Figure 7H). In FLT3-WT cells, p-FLT3 was suppressed to approximately 8% of the DMSO control, indicating near-complete kinase inhibition, whereas FLT3-D835Y and FLT3-F691L cells retained approximately 48% and 72% residual phosphorylation, respectively. Total FLT3 protein was unchanged across all conditions, confirming that Quizartinib does not affect protein stability. Together, these cell-based results provide orthogonal experimental support for the mechanistic hierarchy predicted by the molecular dynamics simulations.

## Discussion

Quizartinib resistance mediated by D835Y and F691L represents a major clinical obstacle in FLT3-targeted therapy, yet the atomic-level mechanisms underlying these resistance phenotypes have remained incompletely characterized. The present MD simulation study provides a mechanistic framework that explains how two structurally distinct mutations converge on a common functional outcome, namely loss of stable type II inhibitor binding, through divergent molecular pathways.

The D835Y mutation confers resistance primarily by destabilizing the inactive DFG-out conformation required for type II inhibitor binding. Substitution of the negatively charged aspartate with an uncharged, bulky tyrosine perturbs a critical electrostatic network: in wild-type FLT3, Asp835 forms a salt bridge with Arg834 (78% occupancy) and hydrogen bonds with the backbone amides of residues 836–838, stabilizing the activation loop in a compact inactive configuration. The introduced tyrosine instead forms hydrogen bonds with Asp829 of the DFG motif (42% of frames), pulling the activation loop toward an active-like configuration, while its larger side chain introduces steric clashes with Leu818 and Phe830 in the inactive state. Free-energy landscape analysis shows that D835Y lowers inactive-state stability by approximately 4 kJ/mol, enabling access to an alternative active-like state populated in 25% of snapshots. Consistent with the MM-GBSA analysis, the resulting affinity reduction (≈5 kcal/mol, ≈5% on the relative MM-GBSA scale) arises from contributions of both van der Waals and electrostatic terms, with the drop in Cys694 hydrogen-bond occupancy (85% to 45%) being particularly consequential given its role as a conserved hot spot for type II inhibitor recognition. These effects propagate allosterically through the kinase domain, affecting the DFG motif, catalytic loop, and helix αC.

The F691L gatekeeper mutation operates through a mechanistically distinct pathway centered on disruption of the hydrophobic pocket. Wild-type Phe691 provides a large aromatic surface for hydrophobic and π–π packing with Quizartinib. Substitution with the smaller leucine creates a 120 Å³ void that eliminates this contribution and destabilizes the surrounding packing, with van der Waals contributions from Leu616, Cys694, and His809 decreasing substantially. The void becomes partially occupied by an average of 2.3 water molecules, introducing entropic penalties that offset enthalpic gains from water-mediated bridging. The enlarged pocket further allows Quizartinib to sample multiple non-productive binding modes, and structural-network analysis shows that F691L isolates the gatekeeper from the cooperative residue network, fragmenting it into disconnected sub-cavities. Notably, although F691 lies more than 20 Å from the activation loop, F691L induces long-range allosteric coupling that also destabilizes the activation-loop region; this explains why the gatekeeper substitution likewise shifts the conformational equilibrium toward active-like states incompatible with type II inhibitor binding and produces the larger MM-GBSA reduction (≈10 kcal/mol, ≈10%) of the two mutants.

Molecular profiling is increasingly used to refine risk stratification and guide treatment selection in hematologic malignancies (Mao et al., 2020, Zhao et al., 2023). While both mutations confer clinically significant resistance, their mechanistic distinctiveness carries important therapeutic implications. D835Y operates primarily through electrostatic and conformational mechanisms, favoring strategies such as type I inhibitors that bind the active DFG-in conformation. This is consistent with clinical observations that midostaurin and gilteritinib retain activity against D835 mutants. Additional approaches include redesigned type II inhibitors forming alternative electrostatic interactions with Asp829, allosteric inhibitors stabilizing the inactive conformation irrespective of activation loop configuration, and covalent inhibitors targeting Cys694. F691L, by contrast, is better addressed through inhibitors with reduced steric bulk capable of productively occupying the enlarged pocket, flexible linker designs that accommodate distorted pocket geometry, or compounds with enhanced hydrophobic potential to compensate for lost gatekeeper contacts. Importantly, neither mutation completely abolishes Quizartinib binding, suggesting that next-generation inhibitors rationally designed to address specific molecular deficits identified by per-residue energy decomposition could restore clinical efficacy. From a precision medicine perspective, mutation-guided inhibitor selection represents a promising strategy worthy of prospective clinical evaluation (Zhao et al., 2019), with type I or allosteric agents preferred for D835 patients and alternative scaffolds optimized for enlarged gatekeeper pockets considered for those harboring F691L. Beyond overcoming target-site resistance, the clinical development of next-generation FLT3 inhibitors should also incorporate cardiovascular safety assessment, given the potential impact of treatment-related cardiovascular complications on long-term outcomes and treatment continuity (Ahmadzadeh et al., 2026).

Several important questions remain for future investigation. The present study focused on single-point mutations D835Y and F691L, whereas clinical samples frequently harbor compound mutations or alternative substitutions such as D835H, D835V, and F691I. Systematic computational analysis of the complete spectrum of clinically observed mutations would provide more comprehensive structure-activity relationships and potentially reveal mutations with divergent mechanisms. Investigation of next-generation FLT3 inhibitors including gilteritinib and crenolanib within the same computational framework would enable direct mechanistic comparison across chemotypes. Extending simulations to multi-microsecond timescales could capture rare conformational transitions between inactive and active states, providing direct observation of activation mechanisms currently inferred from ensemble analyses. The cellular validation reported here (CCK-8 viability and p-FLT3 Western blot in isogenic Ba/F3 lines) confirms the predicted resistance hierarchy; further biophysical validation through isothermal titration calorimetry, hydrogen-deuterium exchange mass spectrometry, X-ray crystallography or cryo-EM, and NMR spectroscopy would provide additional quantitative thermodynamic and structural precision.

The present study offers several methodological strengths. The use of 200-ns explicit solvent MD simulations in triplicate captures conformational dynamics inaccessible to static crystal structure analysis, properly accounting for water-mediated interactions, thermal fluctuations, and entropic contributions. The integrative analytical framework, combining global stability metrics, PCA-based free energy landscapes, MM-GBSA per-residue decomposition, protein-ligand interaction profiling, and binding pocket geometry characterization, provides a multidimensional mechanistic picture that no single analysis could yield alone. Triplicate simulations with statistical correction ensure reproducibility of the reported trends.

Nonetheless, several limitations merit consideration. MM-GBSA does not rigorously account for entropic contributions, and more rigorous alchemical methods such as free-energy perturbation or thermodynamic integration would provide greater quantitative accuracy, albeit at 100- to 1000-fold greater computational cost; accordingly, the MM-GBSA energies reported here should be interpreted as relative, rank-order indicators rather than absolute thermodynamic quantities. Consistent with this, the modest reductions (≈5 and ≈10 kcal/mol) are best read as ordering the three systems (F691L > D835Y > WT) rather than as predictions of absolute affinity changes, and the corresponding cellular IC50 shifts (≈7.9- and ≈20-fold) agree with this ordering. Simulations initiated from a single crystal structure may introduce conformational sampling bias, and enhanced-sampling approaches such as replica-exchange MD or metadynamics could reveal additional low-energy states not accessed within the present timescale. Force-field limitations in describing aromatic interactions and electronic polarization introduce systematic errors, although the qualitative trends and mechanistic interpretations are expected to remain robust. Finally, the cell-based assays establish the direction and ranking of the effects but were performed in an ectopic Ba/F3 system and are not a substitute for quantitative biophysical affinity measurements.

## Conclusions

This computational investigation, using triplicate 200-ns MD simulations, elucidates the molecular mechanisms by which D835Y and F691L confer Quizartinib resistance in AML. On the relative MM-GBSA scale, D835Y reduces the mean binding free energy by approximately 5 kcal/mol (≈5%) through disruption of the activation-loop electrostatic network and hinge hydrogen bonding, with enhanced conformational heterogeneity favoring active-like states. F691L produces a larger reduction of approximately 10 kcal/mol (≈10%), dominated by loss of hydrophobic and π–π gatekeeper contacts together with allosteric effects on the hinge region. Both mutations destabilize the inactive kinase conformation, increase protein flexibility, and fragment the binding-site residue network. CCK-8 and Western blot assays in Ba/F3 cells reproduced the predicted resistance hierarchy (F691L > D835Y > WT), with IC50 shifts (1.2, 9.5, and 24.0 nM for WT, D835Y, and F691L, respectively) and residual p-FLT3 phosphorylation consistent in rank order with the computed reductions. These mechanistic insights provide guidance for the rational design of next-generation FLT3 inhibitors to overcome resistance in FLT3-mutated AML.

## Data Availability Statement

The initial FLT3 kinase domain–Quizartinib crystal structure used as the starting model for the simulations is publicly available from the Protein Data Bank (PDB ID: 4RT7). The input structures, ligand topology and force-field parameters, GROMACS control files, and representative production trajectories generated in this study are available from the corresponding author on reasonable request.

## Author Contributions

DZ: Conceptualization, Methodology, Software, Formal Analysis, Data curation, Writing – original draft. YL: Investigation, Data curation, Visualization, Writing – review and editing. AV: Investigation, Data curation, Writing – review and editing. YM: Formal Analysis, Validation, Data curation, Writing – review and editing. XW: Formal Analysis, Validation, Data curation, Writing – review and editing. BW: Formal Analysis, Validation, Writing – review and editing. YQ: Writing – review and editing. SS: Data curation, Writing – review and editing. XY: Data curation, Writing – review and editing. ZS: Project Administration, Conceptualization, Supervision, Writing – original draft. JN: Conceptualization, Supervision, Writing – review and editing.

