## supplemental figures for "Molecular Mechanisms of Quizartinib Resistance in FLT3 Mutants"

**Divergent Electrostatic and Hydrophobic Mechanisms Underlie Quizartinib Resistance in FLT3 D835Y and F691L Mutant Acute Myeloid Leukemia**

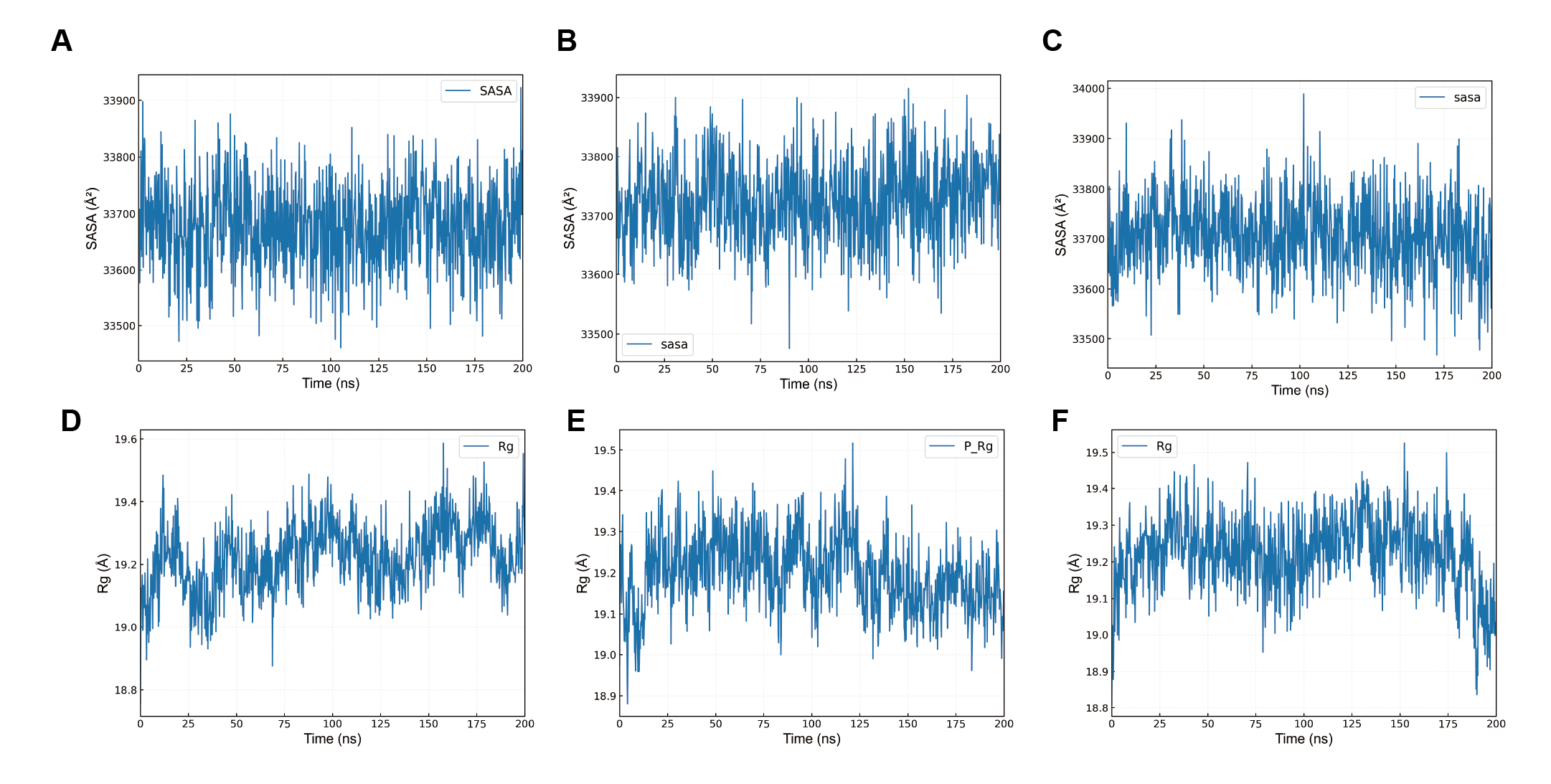

**Figure S1. Dynamics of FLT3 systems during MD. (A–C)** Solvent-accessible surface area (SASA) of FLT3-WT **(A)**, FLT3-D835Y **(B)**, and FLT3-F691L **(C)** complexes. **(D–F)** Radius of gyration (Rg) of FLT3-WT **(D)**, FLT3-D835Y **(E)**, and FLT3-F691L **(F)** complexes.

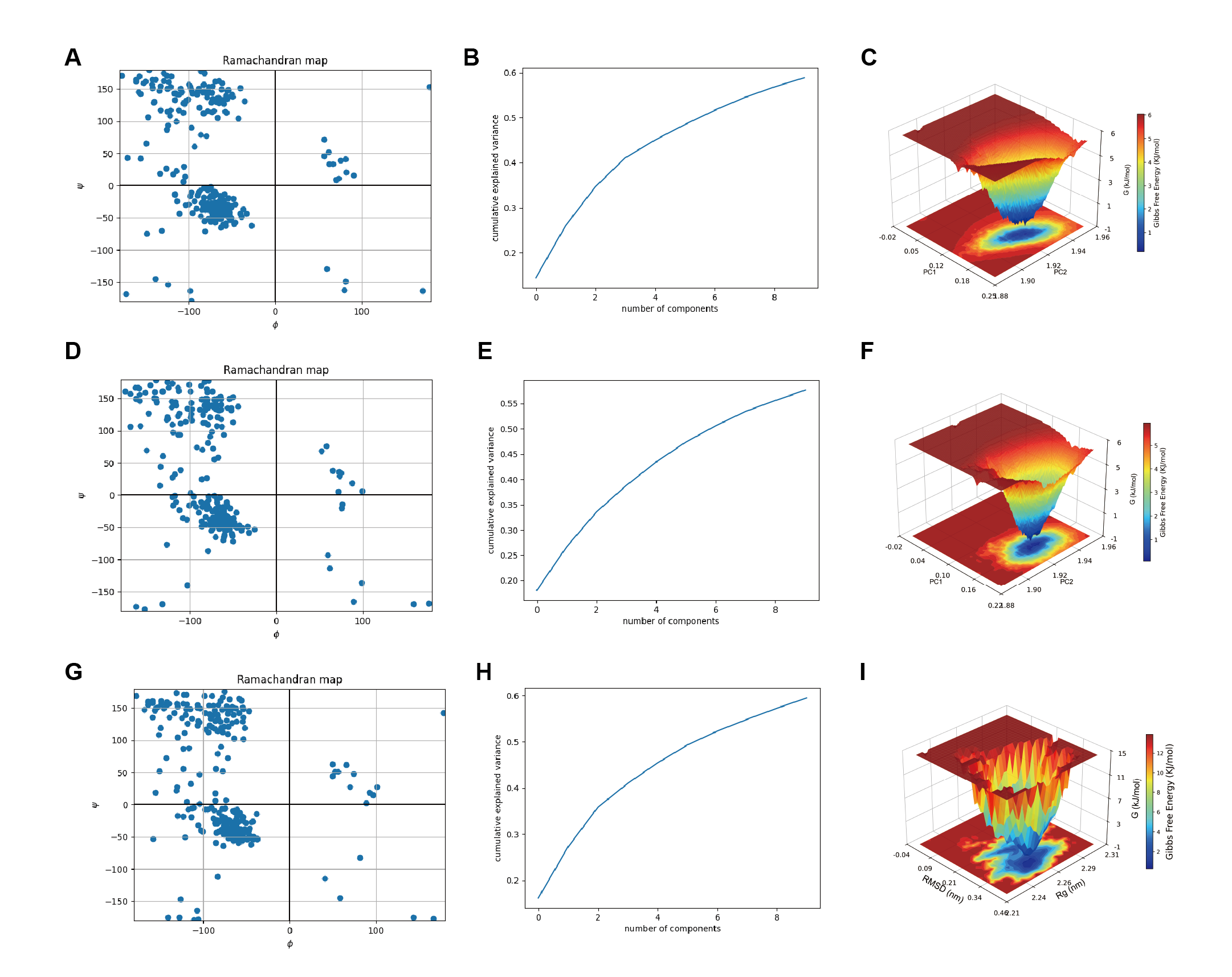

**Figure S2. Conformational dynamics and energy landscapes. (A, D, G)** Ramachandran plots showing backbone dihedral-angle distributions of FLT3-WT **(A)**, FLT3-D835Y **(D)**, and FLT3-F691L **(G)** complexes. **(B, E, H)** Cumulative variance explained by PCA components for FLT3-WT **(B)**, FLT3-D835Y **(E)**, and FLT3-F691L **(H)** complexes. **(C, F, I)** Three-dimensional free-energy surfaces illustrating conformational basins for FLT3-WT **(C)**, FLT3-D835Y **(F)**, and FLT3-F691L **(I)** complexes.

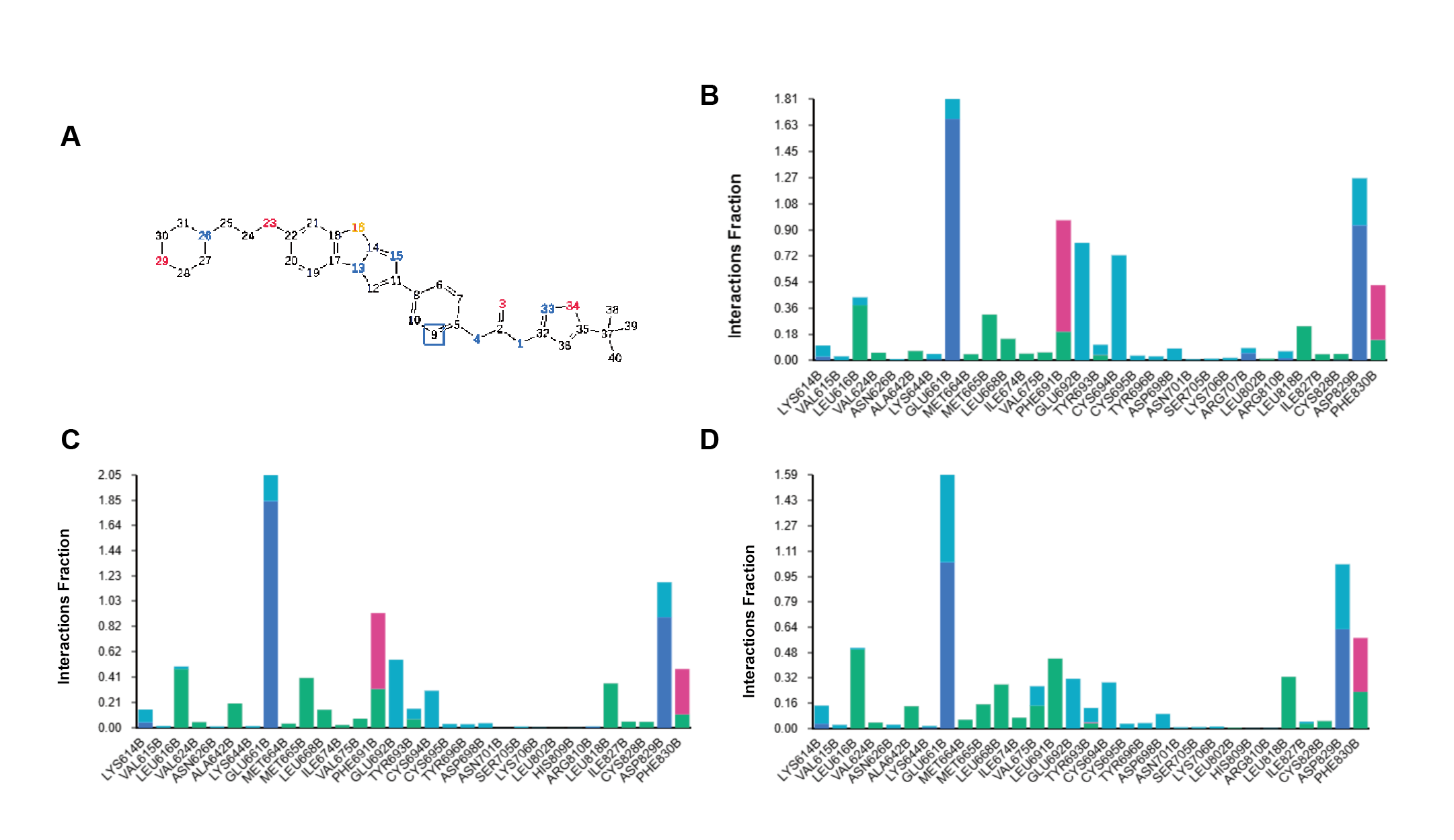

**Figure S3. Molecular interactions in the FLT3–Quizartinib complexes. (A)** Quizartinib atom numbering with red (O), blue (N), and yellow (S) atoms. **(B)** Interaction-frequency histogram of key residues in the FLT3-WT complex. **(C)** Interaction-frequency histogram of key residues in the FLT3-D835Y complex. **(D)** Interaction-frequency histogram of key residues in the FLT3-F691L complex.

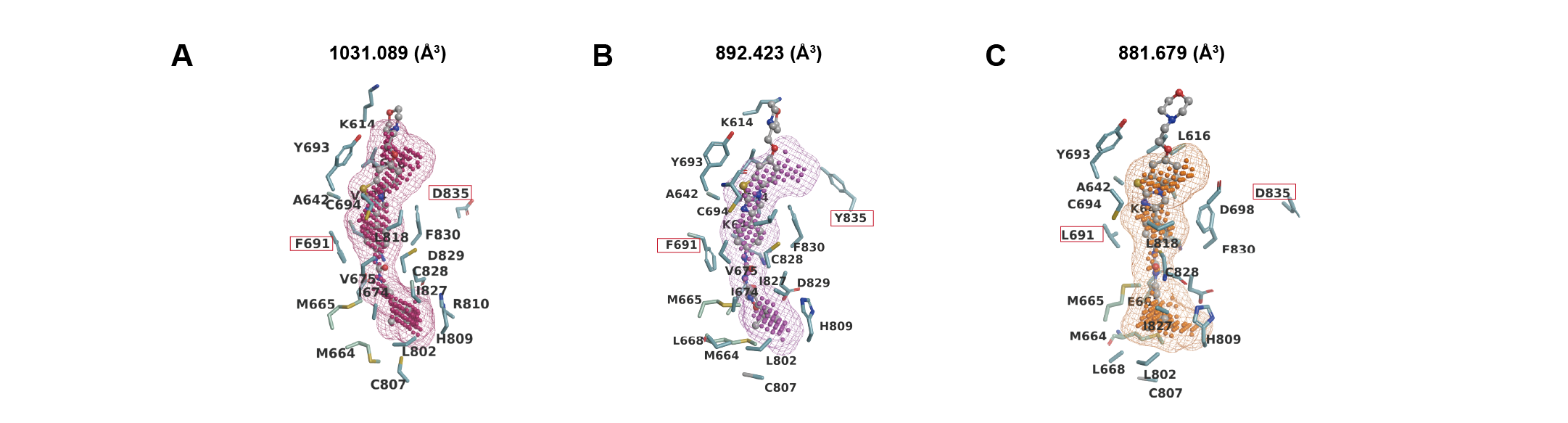

**Figure S4. Binding pocket structural parameters. (A–C)** Residue interaction network contact-area maps of FLT3-WT **(A)**, FLT3-D835Y **(B)**, and FLT3-F691L **(C)** complexes.
